# Bioclimatic modelling of the spread of the West Nile virus in Europe, with a special focus on Ukraine

**DOI:** 10.1101/2024.11.05.622034

**Authors:** S.V. Mezhzherin, V.M. Tytar, I.I. Kozynenko

## Abstract

West Nile virus (WNV) is a re-emerging zoonotic pathogen transmitted by mosquitoes and causes fever and encephalitis in humans, equines, and birds.While all WNV lineages can cause human illness, the lineage WNV2 has gained particular attention in Europe due to its rapid spread and potential for causing severe disease, with large outbreaks happening in summer months.While WNV is not as prevalent in Ukraine as in some other regions of Europe, outbreaks do occur periodically. This is becoming an alarming trend. In this study we focused on the ability of bioclimatic predictors to envisage WNV outbreaks in Europe, making a special emphasis on Ukraine. For this purpose we employ a machine learning methodology for drawing predictions and the SHAP framework from XAI (i.e., eXplainable artificial intelligence) to rank and uncover the most influential WNV drivers.In terms of bioclimatic predictors, the most important for SDM constructions at the European scale was the mean daily air temperature of the coldest quarter and temperature seasonality for Ukraine. Our model suggests that under an impending health threat are areas in the west (excluding the Carpathian highlands) and south of Ukraine.

## Introduction

West Nile virus (WNV), a member of the family *Flaviviridae*, genus *Flavivirus*, is a re-emerging zoonotic pathogen. Currently, West Nile virus (WNV) is considered the most widely spread arbovirus on the planet [1]. WNV is transmitted by mosquitoes and causes fever and encephalitis in humans, equines, and occasionally wild birds. Historically, WNV was reported to be widely endemic in South Africa, and human infections tended to be sporadic, with large epidemics occurring only when unusually high rainfall or hot weather favors breeding of the mosquito vectors. The occurrence of disease has dramatically expanded including the Americas as well as Europe and countries facing the Mediterranean Basin [2]. While lineage 1 of the WNV was the initial introduction to Europe, other lineages, such as lineage 2, have also been introduced and circulated here, contributing to the ongoing spread and evolution of the virus [3]. In Europe, lineage 1 was replaced by lineage 2 in 2013 and currently dominates the transmission [4]. While all WNV lineages can cause human illness, the lineage WNV2 has gained particular attention in Europe due to its rapid spread and potential for causing severe disease, with large outbreaks happening in summer months, particularly in Central and Eastern Europe, however these have only marginally affected Ukraine [5, 6]. In the past ten years a sub-lineage WNV2a which emerged in 2004 has become the dominant lineage [7]. Also outbreaks of WNV2a can have economic consequences, particularly in regions heavily reliant on tourism or agriculture. With regard to health issues in Ukraine, since the start of the full-scale Russian invasion in 2022, millions of people have fled their homes and are displaced inside the country. Such refugee movements can be associated with an increase in infectious disease transmission and are likely to affect zoonotic disease risks.

Even though there have been studies and reports on WNV in Ukraine (for instance, [8]), the availability of genome sequences is relatively limited largely due to budgetary and/or infrastructural challenges. In addition, despite the fact that WNV has been known in the country since the 1970s, official statistics were only launched in 2006 [9].

While WNV is not as prevalent in Ukraine as in some other regions, outbreaks do occur periodically. According to some internet resources, for instance https://hromadske.radio/, as of the beginning of summer 2024, 17 patients have been hospitalized in Kyiv with a clinically confirmed WNV (fever). The specific fever has led to three deaths in the capital. According to the Kyiv City State Administration, among the sick are 10 men and 7 women. All of them lived in different districts of the capital and Cherkasy region.

This is becoming an alarming trend, but the exact causes behind the increased frequency, intensity, and geographic range of WNV outbreaks remain elusive due to the intricate nature of its transmission and the complex factors influencing its spread.

Factors, like temperature and precipitation, geography, and land use, influence WNV spread, and among these, temperature is particularly important in Europe, affecting WNV activity [5]. In this study, we focused on the ability of bioclimatic predictors to envisage WNV outbreaks in Europe, making a special emphasis on Ukraine. For this purpose we employ a machine learning modelling approach for drawing predictions and the SHAP framework from XAI (i.e., eXplainable artificial intelligence) to rank and uncover the most influential WNV drivers [10].

## Materials and methods

Among the lineages found in Europe, WNV2a (a sub-lineage of WNV2) has been predominant, accounting for 82% of all sequences obtained in Europe that have been shared in the public domain [7]. Therefore, we have studied WNV2a in detail using published sequences collected from different host types, with 30% from birds, 30% from mosquitoes, and 40% from humans and other mammals [7]. Faced with limited data for Ukraine, we built species distribution models (SDMs) using the georeferenced locations of these records. Duplicate records were excluded in SAGA GIS (https://saga-gis.sourceforge.io/en/index.html).

The ‘flexsdm’ R (v. 3.3.3) modeling package [11] was used for projecting the potential geographic distribution of WNV2a across Europe, including Ukraine. Spatial block partitioning was used to generate pseudo-absence and background points. Filtering the occurrence data was used to reduce sample bias by randomly removing points where they were dense (oversampling) in the environmental and geographical spaces. The ‘flexsdm’ package offers a wide range of modeling options. Here, we tested out Maximum Entropy (Maxent), one of the most popular SDM modelling methods. Maxent can construct simple to highly complex, nonlinear species–environment relationships using various transformations of variables termed features and represented by a number of feature classes (FC) of which we tested linear, quadratic, product and hinge. To reduce overfitting, Maxent uses a regularization procedure to balance model fit with complexity, by penalizing models based on the magnitude of their coefficients. Tuned models were built using regularization multiplier values ranging from 0 to 4 with increments of 0.5 and all possible FC combinations.

Models were evaluated using the area under the receiver operating characteristic curve (AUC) and the true skill statistic (TSS). AUC scores range from 0 to 1, with 1 for systematically perfect model predictions; AUC values>0.8 are considered to be good to excellent. TSS values range from -1 to +1, with -1 corresponding to systematically wrong predictions and +1 to systematically correct predictions; TSS values>0.5 are considered good to excellent. Because AUC has its drawbacks, we also employed the continuous Boyce index provided by the ‘flexsdm’ package. It is continuous and varies between -1 and +1. Positive values close to 1 indicate a model which present predictions are consistent with the distribution of presences in the evaluation dataset.

SDMs are primarily climate-driven, meaning that the variables used to develop them typically portray climatic factors. This makes sense because climate is a chief driver of environmental suitability and significantly influences the transmission of infectious diseases. Information on bioclimatic parameters was collected as raster layers from the CHELSA climatic data base (https://chelsa-climate.org/) and used for building the anticipated SDMs and checking their performances. Bioclimatic parameters are derived from the monthly temperature and precipitation values in order to generate more biologically meaningful variables. These are often used in species distribution modelling and related ecological modelling techniques. The bioclimatic variables represent annual trends (e.g., mean annual temperature, annual precipitation), seasonality (e.g., annual range in temperature and precipitation) and extreme or limiting environmental factors (e.g., temperature of the coldest and warmest month, and precipitation of the wet and dry quarters).

Commonly used approaches recommend removing correlated predictor variables before modeling to avoid multicollinearity, which affects model projections. The ‘flexsdm’ package offers functions that reduce collinearity in predictors, however in our work they were not employed because the benefits of using all available variables may outweigh the drawbacks of collinearity. Latest research indicates that modelling with correlated climate variables increases accuracy of predictions [12].

In the next step we post-processed the best model results with SHAP [10]. SHAP ranks predictor importance by comparing what a model predicts with and without the predictor for all possible combinations of predictors at every single observation. The predictors are then ranked according to their contribution for each observation and averaged across observations. Importantly, the SHAP method enables us to identify drivers of WNV transmission by region. Another useful item are dependence plots. They show the relationship between a specific predictor and the SHAP values and highlight how changes in the predictor value impact the outcome model. In our case, the R package ‘shap-values’ (https://github.com/pablo14/) in a modified version was used to perform the SHAP analysis.

Maps of habitat suitability (HS) in the GeoTIFF format were processed and visualized in SAGA GIS (https://sourceforge.net/projects/saga-gis/).

Statistical data was analyzed using the PAST software package (https://en.wikipedia.org/wiki/Paleontological_Statistics) and the R environment (https://www.r-project.org/), in particular packages ‘trafo’ (https://doi.org/10.32614/CRAN.package.trafo) and ‘SpatialPack’ (https://doi.org/10.32614/CRAN.package.SpatialPack).

## Results and discussion

From published sources we had at our disposal 301 georeferenced occurrences scattered across countries belonging predominantly to the European Union. Duplicate records were excluded, reducing their number to 156. This number of encounter records is considered more than sufficient to generate reliable SDMs.

The resulting Maxent model (number of filtered occurrences: 73, regularisation multiplier: 1.0, FC combinations: linear×quadratic×hinge) showed high performance with means±standard deviations of AUC and TSS reaching 0.91±0.03 and 0.74±0.11, respectively, and a continuous Boyce index of 0.92±0.06.

In terms of bioclimatic predictors, the top 5 most important for SDM constructions at the European scale were: mean daily air temperatures of the coldest quarter (bio11), mean daily minimum air temperature of the coldest month (bio6), temperature seasonality (bio4), mean daily air temperatures of the wettest quarter (bio8) and isothermality (bio3) (Fig.1A). Obviously, amongst these, bio11 plays an outstanding role in shaping the spread of the WNV, which strongly correlates with the final SDM (the corrected Pearson’s correlation for spatial autocorrelation amounts to 0.84, p<0.05). Most likely this role is mediated through the mosquito vector, thus highlighting the importance of understanding the overwintering ecology of the insect species, in particular *Culex pipiens pipiens* s.l. Typically, transmission cycles of mosquito-borne pathogens are suspended during the winter months. Prior to this, many female mosquitoes in late summer prepare for diapause by accumulating fat reserves. They then migrate to hibernation sites, commonly basements and other underground constructions, where they usually remain until spring. Temperatures between 2 and 6°C have been identified as optimal for overwintering adults [13]. The corresponding dependence plot gained through the SHAP analysis showed an optimum of 2.85°C. Engagingly, this figure was reached by applying a totally dissimilar approach. Furthermore, temperatures lower than 0°C lead to death after several days and higher temperatures increase the metabolic rates and therefore precariously deplete the fat reserves of the mosquito females. Accordingly, under unfavorable winter conditions temperatures can be considered an important factor that has the potential to limit the geographic ranges of the mosquito. Also temperature-dependent is the extrinsic incubation period. Essentially, this is the period during which the mosquito becomes capable of transmitting the disease to another host. For the mosquito species that can transmit WNV this is typically 16-25 days. Warmer temperatures generally shorten this period. Conversely, colder temperatures have an opposite effect. This is why WNV outbreaks tend to occur in warmer regions during the summer months where air temperatures of the coldest quarter on average are higher (mean daily mean air temperatures of the warmest and coldest quarters of the year are positively correlated: accounting for spatial autocorrelation, the corrected Pearson’s correlation is estimated as 0.58, p<0.05). To this end, warmer winters are likely to improve the survival rates of mosquito overwintering populations, potentially leading to larger populations in the following year. Earlier it was shown that high enough temperatures in the month May have an important impact on WNV transmission dynamics throughout the season [14].

**Fig. 1.**
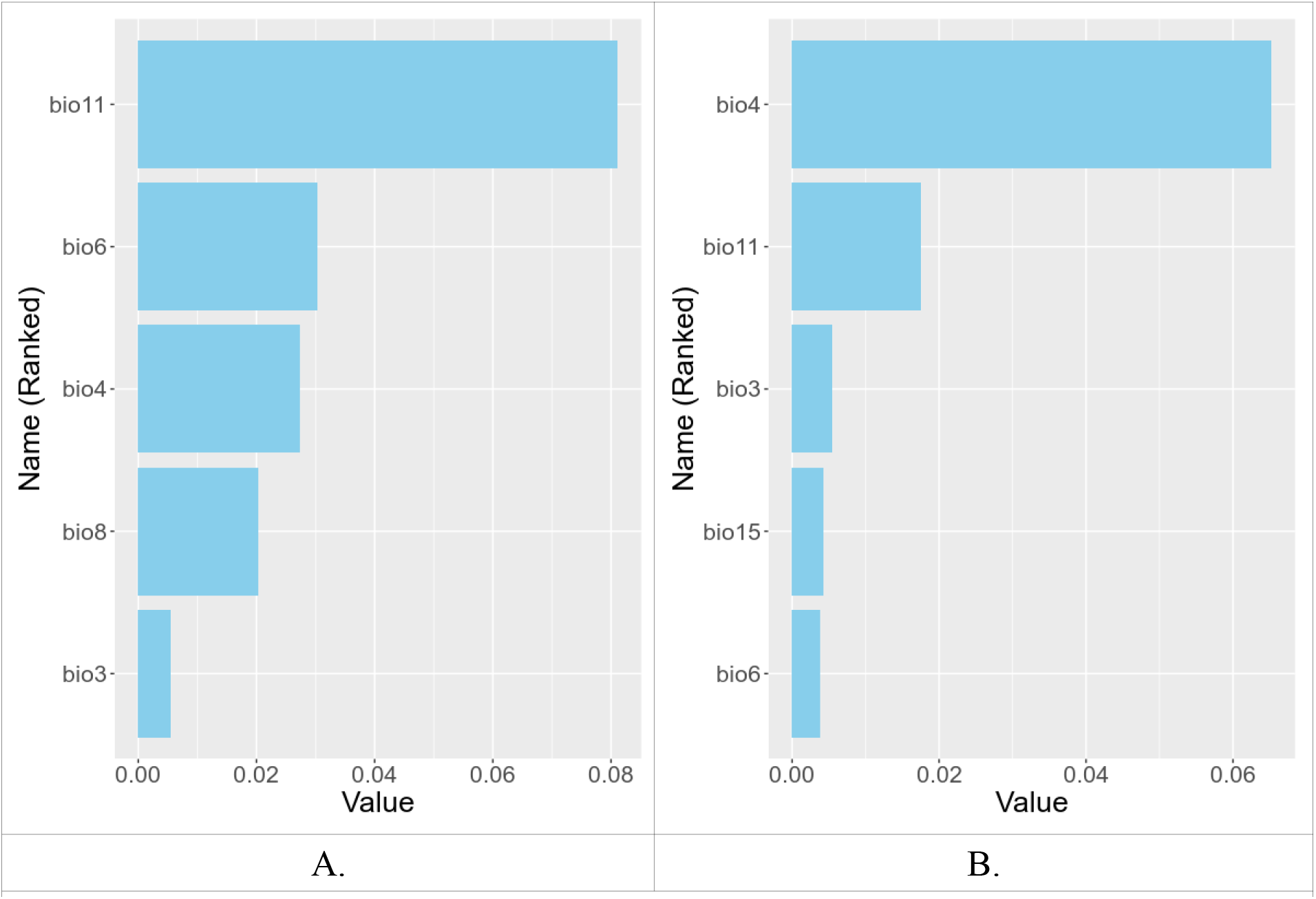
Summary barplots of the average absolute SHAP values of the 5 most contributing to the model variables. The y-axis represents variables used in the study (see text). The *x*-axis represents the corresponding SHAP value. A: Europe, B: Ukraine.

Also warmer spring temperatures may allow mosquitoes to emerge earlier, extending the transmission season. In addition, rising temperatures can create new suitable habitats for mosquito breeding, particularly in regions previously too cold. Indeed, according to the A1B climate change scenario (https://www.ipcc.ch/), mean daily air temperatures of the coldest quarter by 2030 will increase by almost 1.6°C. Finally, warmer winters allow birds, where WNV primarily circulates, to migrate earlier and establish territories sooner [15], although regional variations play significant roles.

As for the territory of Ukraine, the top 5 most important predictors of WNV occurrence are: temperature seasonality (bio4), mean daily air temperatures of the coldest quarter (bio11), isothermality (bio3), precipitation seasonality (bio15), mean daily minimum air temperature of the coldest month (bio6). This time temperature seasonality (bio4) is notably put in the foreground and only then followed by bio11. However, both are interconnected and between them stands a moderate negative corrected Pearson’s correlation of -0.51 (p<0.05), meaning temperature seasonality is on average greater in areas where winter temperatures are low and vice versa.

SDMs are valuable tools for predicting the geographic distribution of species, including diseases and their vectors. When applied, these models often employ thresholds to delineate areas of high and low risk. In this respect, because of the hazardous character of West Nile fever, we gave preference to the minimum training presence (MTP), a parameter commonly used in SDMs to ensure that a sufficient number of occurrence records are present within each environmental cell or grid. This helps to prevent the model from overfitting to areas with high occurrence density and improves the generalization of predictions to areas with fewer or no records. The corresponding SDM for the WNV in Ukraine and the potential disease risk area are presented in Fig.2.

**Fig. 2.**
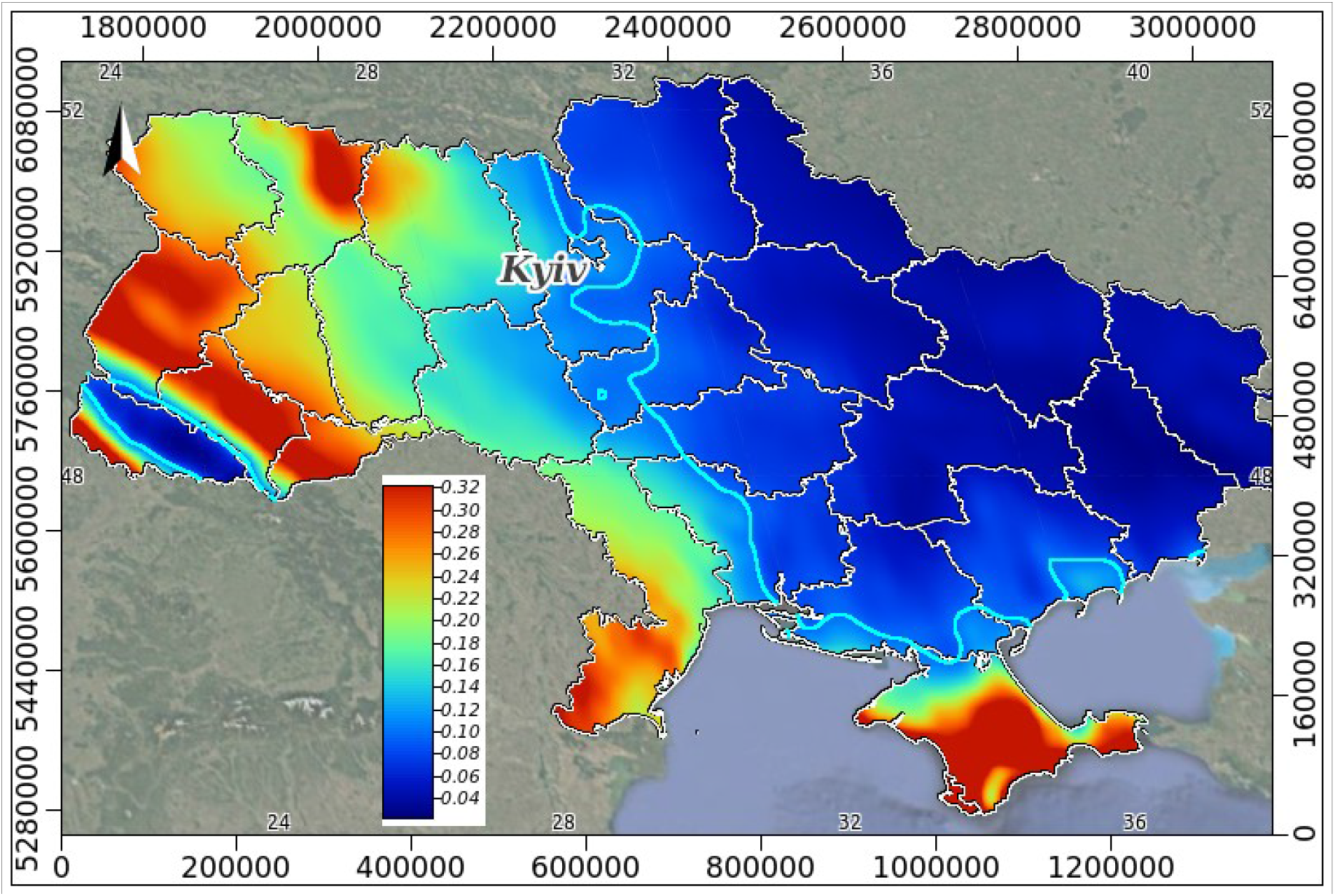
A habitat suitability (HS) map in Mollweide’s equal-area pseudocylindrical projection for the West Nile virus in Ukraine; the legend shows potential HS ranging from high (red) to low (navy blue); the azure line represents the minimum training presence (MTP) threshold of 0.096.

Ranked by maximum values of HS arbitrarily exceeding 0.3, the most affected regions in the country could be: Crimea > Sevastopol’ > Transcarpathia > L’viv > Chernivtsi > Ivano-Frankivs’k > Rivne > Odessa > Ternopil’ > Zhytomyr > Volyn’. Generally speaking, under an impending health threat are areas in the west (excluding the Carpathian highlands) and south of Ukraine. Therefore, a question raises of how safe are these regions in terms of disease transmission. Particularly this concerns the west where millions of people have found an escape from the war, but according to the constructed model are put at a higher risk of infection. Hopefully, our findings could substantially help health officials in managing and monitoring WNV distributions. Curiously, our SDM covers the capital, Kyiv, and a portion of Cherkasy region where local incidents of WN fever have been recorded this year.

## Conclusion

WNV is an emerging mosquito-borne pathogen in Europe where it represents a new public health threat. WNV was considered a minor risk for humans because of its sporadic occurrence. However, over the last two decades there have been repeated local outbreaks across various regions worldwide, such as Europe, including Ukraine. Using only bioclimatic features, our modelling approach indicates a strong dependence of WNV occurrence on winter temperatures. Rising temperatures are likely increasing the transmission potential of mosquito-borne diseases in Europe, by affecting the geographic spread, abundance, survival and feeding activity of vector species and benefiting pathogen development in infected vectors. Overall, the design and implementation of preparedness plans for the prevention of cases in humans is an urgent issue, especially under current wartime circumstances.

## Notes

### Competing Interest Statement

The authors have declared no competing interest.

## REFERENCES

1. Costa E.A., Bayeux J.J.M., Silva A.S.G. et al. Epidemiological surveillance of West Nile virus in the world and Brazil: relevance of equine surveillance in the context of “One Health”. Braz. J. Vet. Res. Anim. Sci. [Internet]. 2019, 56, 4, e164335. https://www.revistas.usp.br/bjvras/article/view/164335

2. Zellar H.G., Schuffenecker I. West Nile virus: An overview of its spread in Europe and the Mediterranean Basin in contrast to its spread in the Americas. Eur. J. Clin. MIcrobiol. Infect. Dis. 2004, 23, 3. p. 147–156. DOI: 10.1007/s10096-003-1085-1

3. Koch R.T., Erazo D., Folly A.J. et al. Genomic epidemiology of West Nile virus in Europe. One Health. 2023, 18, e100664. DOI: 10.1016/j.onehlt.2023.100664

4. Calzolari M., Monaco F., Montarsi F. et al. New incursions of West Nile virus lineage 2 in Italy in 2013: the value of the entomological surveillance as early warning system. Vet. Ital. 2013, 49, 3. P. 315–319. DOI: 10.1016/j.onehlt.2023.100664

5. Paz S., Semenza J.C.. Environmental drivers of West Nile fever epidemiology in Europe and Western Asia--a review. Int. J. Environ. Res. Public Health. 2013, 10, 8. P. 3543–3562. DOI: 10.3390/ijerph10083543

6. Napp S., PetriĆ D., Busquets N. West Nile virus and other mosquito-borne viruses present in Eastern Europe. Pathog. Glob. Health. 2018, 112, 5. P.:233–248. DOI: 10.1080/20477724.2018.1483567

7. Lu L., Zhang F., Oude Munnink B.B. et al. West Nile virus spread in Europe: phylogeographic pattern analysis and key drivers. PLoS Pathog. 2024, 20, 1. e1011880. DOI: 10.1371/journal.ppat.1011880

8. Kotelevska T. M., Pryimenko N. O., Dubynska H. M. et al. West Nile Fever in the central part of Ukraine. Medicni perspektivi. 2020. 25, 3. P. 204–210. DOI: 10.26641/2307-0404.2020.3.214874

9. Yushchenko A., et al. Ecological and epidemiological aspects of West Nile virus in Ukraine. Acta Scientific Microbiology. 2020, 3, 5. P. 31–36. DOI: 10.31080/ASMI.2020.03.0584

10. Farooq Z., Rocklöv J., Wallin J. et al. Artificial intelligence to predict West Nile virus outbreaks with eco-climatic drivers. The Lancet Regional Health - Europe. 2022, 17, e100370. DOI: 10.1016/j.lanepe.2022.100370

11. Velazco S.J.E., Rose M.B., Andrade A.F.A. et al. flexsdm: An R package for supporting a comprehensive and flexible species distribution modelling workflow. Methods in Ecology and Evolution, 2022 13, 8. P. 1661–1669. DOI: 10.1111/2041-210X.13874

12. Hanberry B. B. Practical guide for retaining correlated climate variables and unthinned samples in species distribution modeling, using random forests. Ecological Informatics. 2023, 79, 1, e102406. DOI: 10.1016/j.ecoinf.2023.102406

13. Rozsypal J., Moos M., Rudolf I. et al. Do energy reserves and cold hardiness limit winter survival of Culex pipiens? Comp. Biochem. Physiol. A Mol. Integr. Physiol. 2021, 255, e110912. DOI: 10.1016/j.cbpa.2021.110912.

14. Angelou A., Kioutsioukis I., Stilianakis N.I. A climate-dependent spatial epidemiological model for the transmission risk of West Nile virus at local scale. One Health. 2021, 13, e100330. DOI: 10.1016/j.onehlt.2021.100330.

15. Both C., Visser M.E. The effect of climate change on the correlation between avian life-history traits. Global Change Biology. 2005, 11, 10. P. 1606–1613. DOI: 10.1111/j.1365-2486.2005.01038.x

